# Membrane-tethered cadherin substrates reveal dynamic and local shifts in actin network architecture during adherens junction formation

**DOI:** 10.1101/2024.09.09.611904

**Authors:** Sayantika Ghosh, John James, Badeer Hassan Ummat, Helena Coker, Marco Fritzsche, Darius Vasco Köster

## Abstract

Adherens junctions (AJs) are E-cadherin-based adhesions at cell-cell contacts that connect the actin cytoskeleton of epithelial cells. Formation and maturation of these junctions is important in development, e.g. for the generation of epithelial tissues, and loss of adherens junctions is linked to metastasis in cancer. It is well established that AJ formation is a mechano-sensitive process involving cis and trans clustering of E-cadherin and the mechanotransducive activation of α-catenin that connects E-cadherin with the actin cytoskeleton. However, how mobility of E-cadherin in the cell membrane and their local density influences actin polymerisation is less well understood due to limitations in controlling physical properties of cell membranes and performing high-resolution microscopy in model organisms or cell-monolayers. Here we have created a biomimetic system enabling super-resolution microscopy of AJs by placing MCF7 cells, labelled with fluorescent actin, E-cadherin and α-catenin, on fluid supported lipid bilayers (SLB) containing the extracellular domain of E-cadherin. We found that MCF7 cells were able to attach and spread on these substrates, recruiting E-cadherin and α-catenin to form AJs that can mature and are mobile. Interestingly, we found that depending on E-cadherin mobility within the SLB, distinct types of actin architecture emerge. Low mobility substrates support formin-based linear actin polymerisation while high mobility substrates support Arp2/3-based branched actin network polymerisation. These polymerising actin structures are spatially confined to regions of low E-cadherin density suggesting they may play a role in AJ repair. Following how these actin structures at the cell-SLB interface evolve over time indicates regions of both linear and branched actin being present at mature cell-cell contacts.

## Introduction

Cells in epithelial tissues are held together by adherens junctions (AJs) formed by E-cadherin interactions at the apical pole of the cell. These structures serve as mechanical anchors and contact points for biochemical and mechanical signalling. Mechano-signalling from AJs is essential to regulate cellular processes like migration and proliferation (Friedl and Mayor 2017). The mechanical forces generated and sensed by these structures depend on the architecture of the underlying actin cytoskeleton (Charras and Yap 2018; Arslan et al. 2021).

AJs are formed when the extracellular domains of E-cadherin from adjacent cells interact with each other. The intracellular domains of E-cadherin can form a complex with β-catenin and the α-catenin head domain, called the cadherin-catenin-complex (CCC) which represents the structural unit of an AJ and acts as a mechano-sensitive link to the actin cytoskeleton (Mège and Ishiyama 2017, James, Winn, et al. 2025). The force-dependent recruitment of adaptor proteins, such as vinculin, to the CCC, such as vinculin, to the CCC (Yao et al. 2014) promotes clustering of E-cadherin via their extracellular domains, by both cis-clustering along the same plasma membrane and trans-clustering of E-cadherins on the membranes of adjacent cells (Thompson et al. 2021). The classical view of a mature cell-cell contact has linear actin bundles stretched taut between AJs parallel to the cell contact (Cavey and Lecuit 2009). The cell membranes are held together by extracellular E-cadherin interactions and the tension across these actin bundles. A combination of cadherins being internalised in vesicles and flowing in from the surrounding membrane, along with the recruitment of actin modulating proteins constantly replenishes and maintains these AJs (Reynolds 2010).

Actin polymerisation in cells is governed by actin nucleation factors like formins and Arp2/3 which give rise to different actin architectures within the cell (Fritzsche et al. 2013). Formins are a family of proteins that promote nucleation of linear actin filaments (Oosterheert et al. 2024). Arp2/3 is a protein complex that binds existing actin filaments and mimics an actin trimer to create a branch by allowing for linear polymerisation at a 70° angle from the mother filament (Papalazarou and Machesky 2021). Linear actin nucleators and elongation factors such as the formins, Formin1, mDia1 and FMNL2 as well as mENA/Vasp have been reported to be present at AJs (Kobielak et al. 2004; Acharya et al. 2017; Grikscheit et al. 2015; Oldenburg et al. 2015). The branched actin network nucleator Arp2/3 along with its activators RAC1, WAVE, IQGAP2 and cortactin are also recruited to AJs (Braga et al. 1997; Han et al. 2014; Noritake et al. 2004).

Although reported decades ago (Vasioukhin et al. 2000), the effects and regulation of actin polymerisation at AJs are still not entirely clear. The initial study suggested that polymerisation from AJs sustained the linear bundles of actin present at mature cell-cell contacts. Branched actin polymerisation also plays an important role in newly forming cell-cell contacts (Lee et al. 2016), with branched actin networks being remodelled into linear bundles by debranching proteins like Coronin1B (Michael et al. 2016) and actin bundling proteins like myosin (NM2A & NM2B) and eplin (Abe and Takeichi 2008; Yu-Kemp et al. 2021). More recent studies have found branched actin networks at mature cell-cell contacts that could function as a repair mechanism for disassembled AJs (Li et al. 2021; 2020). Since AJs are constantly maintained structures, it is unsurprising that formation and repair (or re-formation) may be governed by the same mechanism (James, Winn, et al. 2025). Indeed, an increase in branched actin polymerisation at AJs has been consistently correlated with increased junction integrity (Senju et al. 2023; Moztarzadeh et al. 2024; James, Fokin, et al. 2025).

Since AJs form between cells at the apical pole of epithelial monolayers, capturing the dynamics of the molecular components of AJs along the cell-cell contact plane using live cell imaging has proven difficult. To overcome these drawbacks, several studies have imaged cells adhering onto E-cadherins immobilised on coverslips (Bertocchi et al. 2017; Schimmel and Noordstra 2023; Collins et al. 2017), but these systems offer limited insights since the lateral mobility of E-cadherin and its clustering cannot be reproduced here. Alternatively, E-cadherin extracellular domain (E-cadECD) tethered to supported lipid bilayers formed on glass coverslips incorporate these parameters and is ideally suited to image AJs at the cell-SLB interface using total internal reflection microscopy (TIRF) (Biswas et al. 2015; 2016; Arslan et al. 2024).

Using this system. we study the organisation of the actin network, E-cadherin and α-catenin in MCF-7 cells, a cell line derived from breast cancer tissue (Noordstra et al. 2023). Using total TIRF and TIRF-structural illumination microscopy (SIM) we follow protein dynamics during AJ formation and maturation up to five hours post adhesion. We were able to recapitulate previously reported characteristics of AJs to confirm the relevance of our model system and discovered that E-cadherin mobility and density direct the formation of regions at the cell-SLB interface that are dominated by either Arp2/3 or formin-driven actin polymerisation. Our results indicate that cells can sense ligand mobility on their neighbours to direct actin polymerisation at AJs.

## Results

### MCF7 cells form adherens junctions on SLBs containing E-cadECD

MCF7 E-cadherin-GFP knock in cells expressing LifeAct-TagRFPT and iRFP670-α-catenin (Noordstra et al. 2023) were plated on supported lipid bilayers (98% DOPC, 2% NiNTA-DGS) decorated with His_6_-E-cad-ECD (Fig. 1a). TIRF imaging revealed that MCF7 cells can attach and spread on these substrates (Supp Fig. 1 b, Supp Video 1). The cell spreading area increased during the first 60 min and then plateaued at 69 (±17) µm^2^, indicating that cell spreading reaches a steady state within an hour (Supp Fig 1b). Cell spreading is driven by actin rich protrusions that grow from the edge of the cell along the SLB. Although the protrusion area decreased over time it persisted once the cell reached maximum spreading (Supp Fig. 1 f-h). We observed that actin, E-cadherin and α-catenin were recruited to the cell-SLB interface showing the formation of AJs (Supp Fig. 1 c-e). Interestingly, the recruitment of AJ proteins continued after the end of cell spreading suggesting further AJ formation and maturation at later time points.

**Fig 1:**
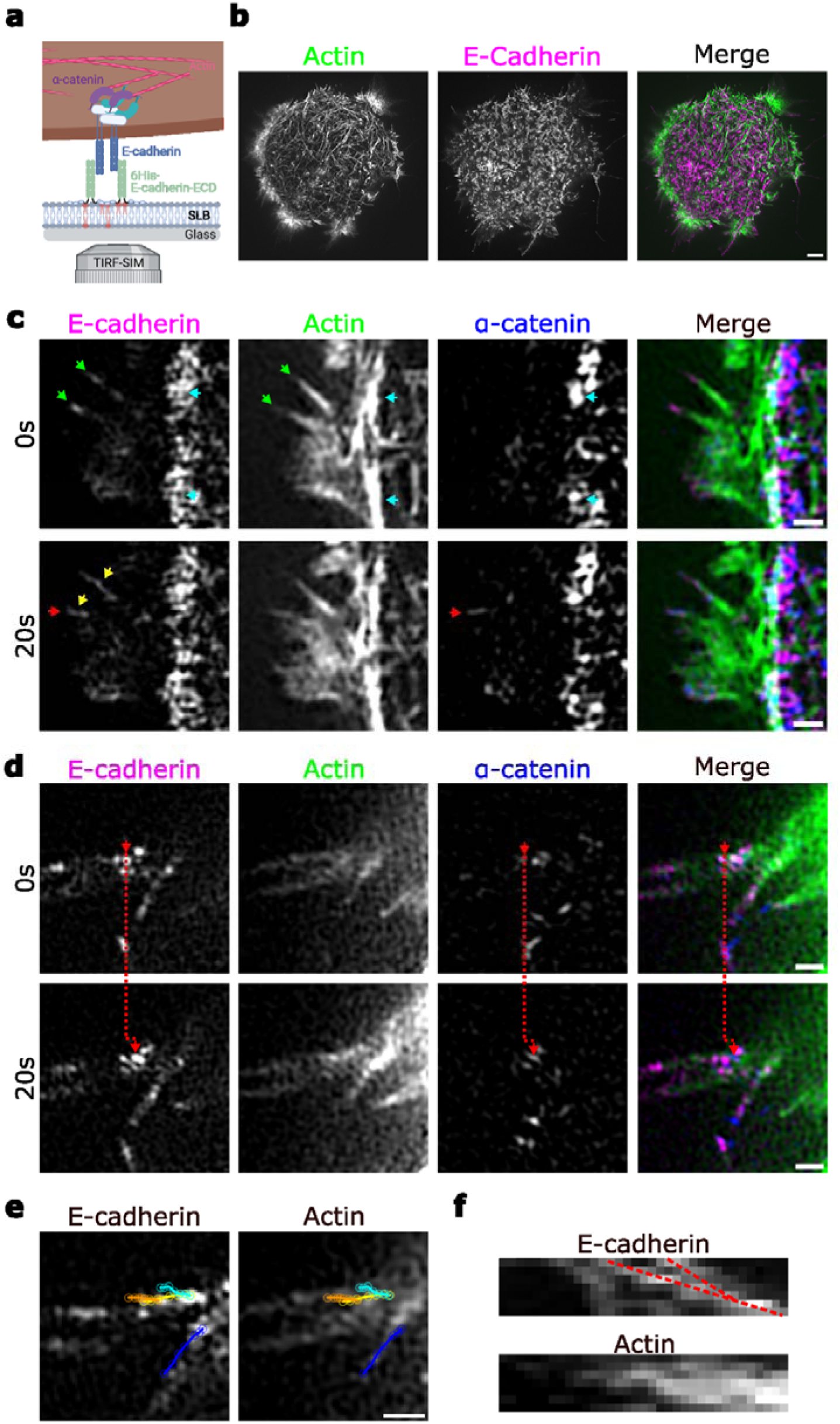
MCF7 cells form mobile AJs on SLBs containing E-cadherinECD. **a** MCF7 E-cadherin-GFP knock in cells expressing LifeAct-TagRFPT and iRFP670-α-catenin were plated on SLBs decorated with His_6_-E-cadECD **b** Sample TIRF-SIM image of a whole cell expressing GFP-E-cadherin and LifeAct-TagRFPT. Scale Bar=2µm **c** TIRF images of MCF7 cells expressing GFP-cadherin, LifeAct-TagRFPT and α-catenin. Green arrows show E-cadherin recruited to ends of Actin filaments. Yellow arrows show Cadherin staining lengthening along the filament. Red arrow shows α-catenin recruited to E-cadherin enriched structures. Cyan arrows show E-cadherin, α-catenin and actin colocalised in mature AJs. **d** E-cadherin and α-catenin puncta move over time **e** Tracks of E-cadherin puncta moving along actin filament. **f** Kymographs of E-cadherin and actin showing the movement of and clustering of E-cadherin puncta. All scale bars are 0.5µm.

To obtain super-resolution images of AJs at different stages of maturation, we turned to TIRF-SIM (Fig 1b). At the edge of protrusions, we found E-cadherin puncta at the tips of actin bundles that are stable over tens of seconds but no α-catenin (Fig 1c). Since TIRF-SIM only excites fluorophores within the first ∼100 nm above the surface of the coverslip, these structures are presumably points of attachment to the E-cadECD-SLB substrate. Following these early adhesions over time revealed their elongation indicating further clustering of E-cadherin along actin protrusions as well as the recruitment of α-catenin (red arrows), which are hallmarks of AJ formation. These AJs move over time along the protrusions (Fig 1d). Single particle tracking showed that AJs move along actin filaments from the tip of the protrusion to the base leading to further clustering (Fig 1e, f, Supp Video 2). Concomitantly, at the base of the protrusion, we see an enrichment of mature AJs indicated by an increase in actin, E-cadherin and α-catenin (Fig 1c, cyan arrows). This enrichment leads to the formation of an actin ring at the perimeter of the cell (Supp Fig 2 a, b) as reported previously (Arslan et al. 2024). Thus, our cell-SLB system recapitulates key features seen in physiological AJs (Noordstra et al. 2023).

**Fig 2:**
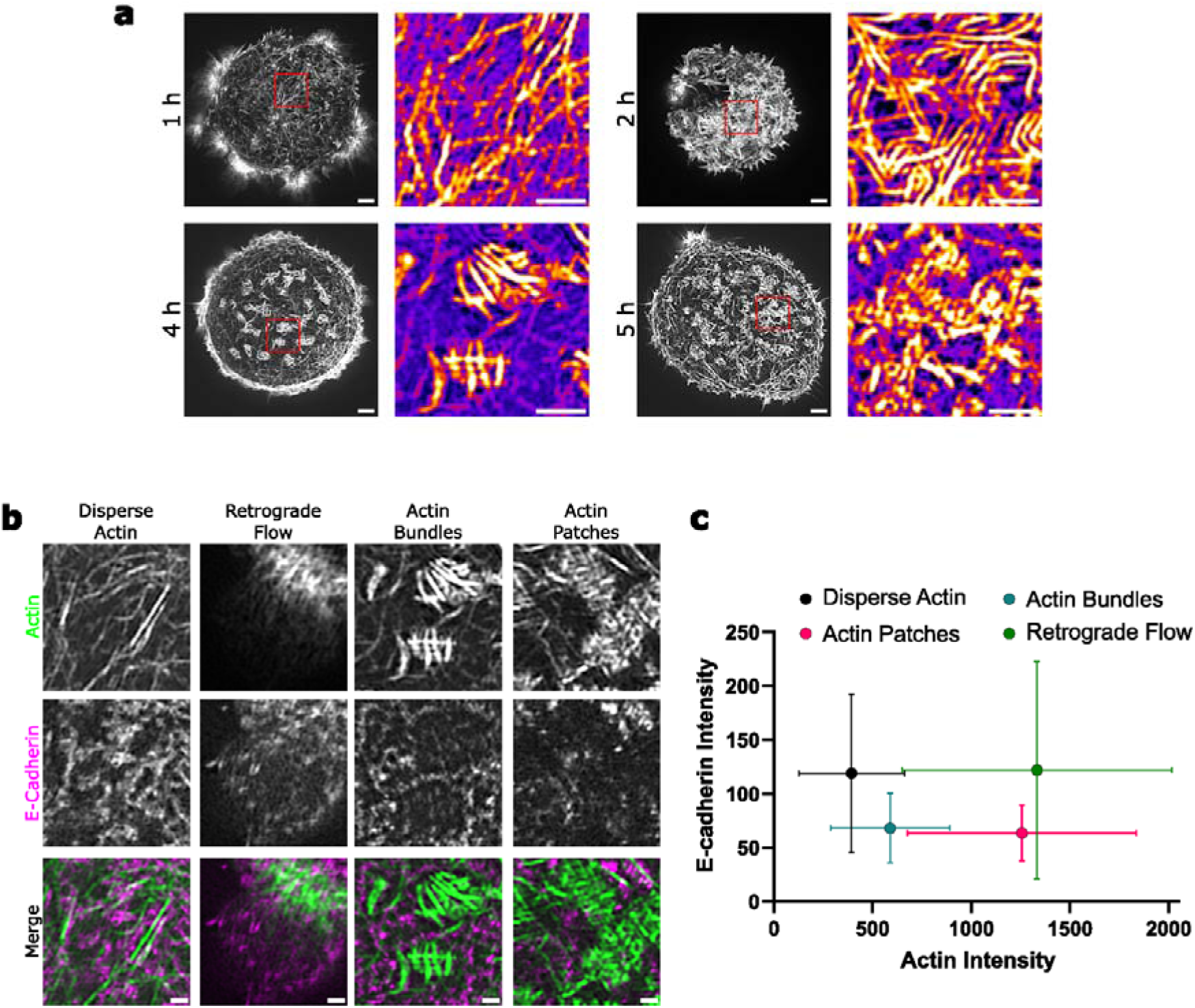
**a** TIRF-SIM images of actin in MCF7 cells imaged at different times post-seeding. Scale Bar = 2µm **b** TIRF-SIM images showing E-cadherin recruitment at regions of different actin architectures Scale Bar = 0.5µm **c** Average intensities of LifeAct-TagRFPT and GFP-E-cadherin in regions of different actin architecture, made by binning 8×8 pixels from TIRF-SIM images to make 250nm windows). Mean±SD plotted.

### Actin polymerisation leads to dynamic actin architectures at the cell-SLB interface

Next, we looked at the cell interior as a model for mature cell-cell contacts that have been in contact with the SLB for up to 5 hours. We found that the actin architecture at the interface is dynamic and evolves over long time scales (Fig 2a). The actin architectures we found are different from those in cells plated on glass coverslips coated with immobile E-cadherin substrates (Collins et al. 2017), with the absence of stress fibres being the most striking observation.

We identified four types of actin structures at the cell-SLB interface - disperse actin, retrograde flow, actin bundles and actin patches (Fig. 2b, Supp Video 3). Retrograde flow is found at the cell edge and in protrusions, like reported of cells plated on substrates with immobilised E-cadherin or extra-cellular matrix proteins. The E-cadherin distribution in this region is characterised by the E-cadherin puncta at the tips of protrusions and mature AJs at the base. Disperse actin are long actin filaments spanning the cell-SLB interface and that are decorated by E-cadherin puncta, while actin bundles and patches seem to appear in spatially delimited regions of low E-cadherin. Both static bundles and disperse actin seem to be distinct from stress fibres observed in cells on immobilised e-cadherin and extra-cellular matrix, as they do not show the characteristic low curvature expected for fibres under tension (Supp. Fig. 2e). Actin patches are characterised by high intensities of actin with the absence of long linear structures.

Plotting the intensities of actin and E-cadherin within 250 nm windows in each region revealed that the four types of actin structures show distinct correlations (Fig. 2e, Supp Fig. 2c, d). Regions of retrograde flow were characterised by high levels of E-cadherin and actin Interestingly, both actin bundles and actin patches are formed in regions of decreased E-cadherin intensity with actin patches showing higher levels of actin.

We next performed time-lapse imaging of these structures to study their dynamics. Disperse actin seemed to move indiscriminately within 5 minutes, and actin bundles and patches showed internal dynamics that remained localised to specific regions (Fig 3a). By quantifying the rate of photobleaching in different actin structures we assessed quantitatively how quickly they remodel and incorporate new fluorescent actin probes (Fig. 3b, c). Disperse actin has the highest photobleaching rate, ∼1.7 times faster than actin bundles indicating that actin bundles are actively maintained by actin polymerisation. Indeed, imaging at shorter time-intervals and tracking the ends of these linear structures revealed that actin bundles elongate over time (Fig 3d, Supp Video 4).

**Fig 3:**
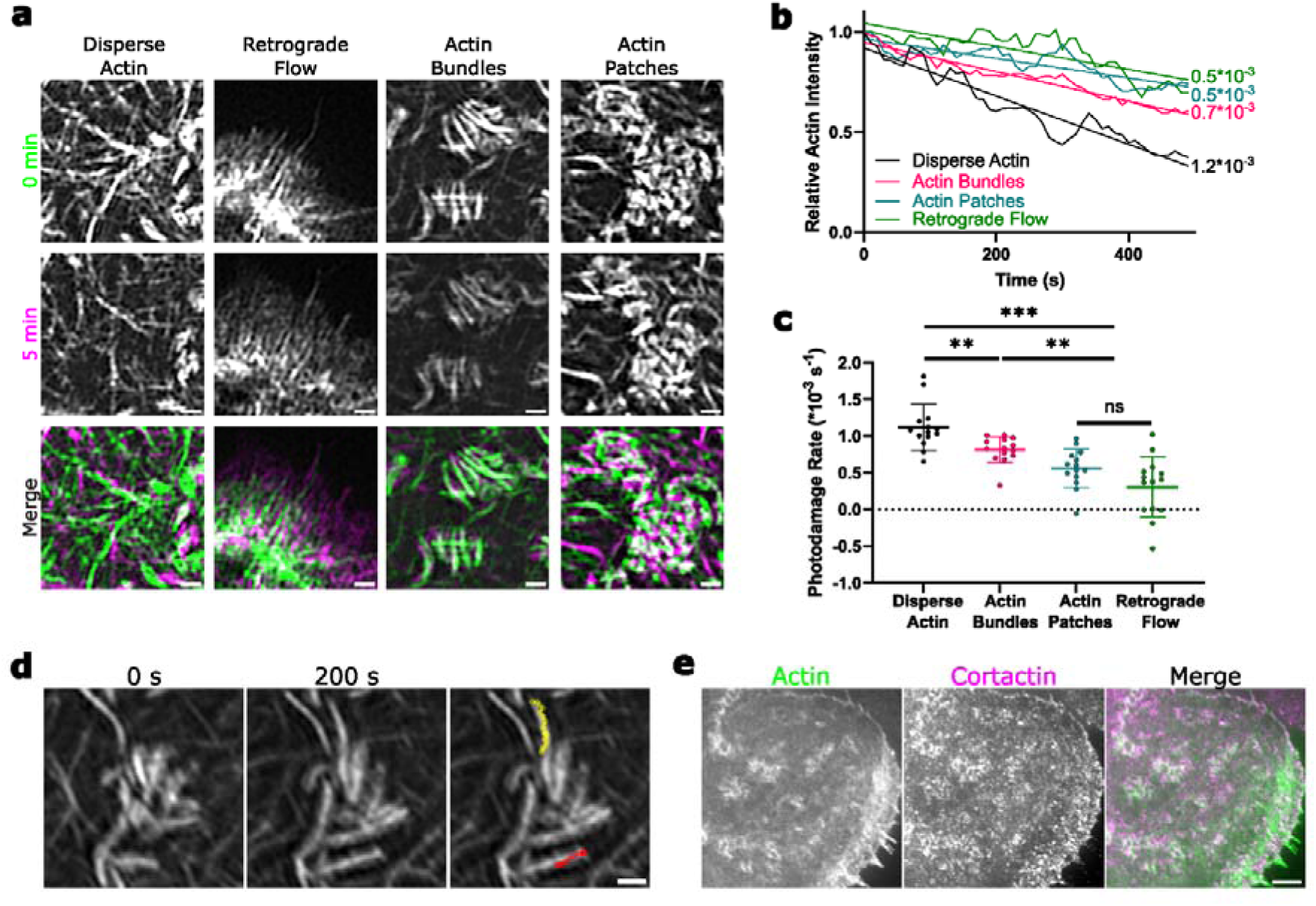
**a** TIRF-SIM images of different actin architectures before and after imaging for 5 mins at an interval of 10 s per image. Scale Bar = 0.5µm **b** Representative curves of decrease in actin signal intensity over time in different actin architectures. **c** Rate of decrease in actin intensity in regions with different actin architectures. Mean ± SD plotted n=10 from N=2. **d** TIRF-SIM images showing extension of actin bundles (red and yellow lines shown track of the bundle tip over time) Scale Bar = 0.5µm **e** Confocal images of MCF7 cells expressing LifeAct-GFP stained with antibodies against the branched actin marker, cortactin.

The lowest photobleaching rates are found in regions of retrograde flow and actin patches (2.4 times slower than disperse actin), suggesting a faster mode of actin filament renewal and remodelling than in actin bundles (Fig. 3 b, c). We assumed that these structures are maintained by branched actin polymerisation, since actin branching amplifies the number of barbed ends available for polymerisation (Achard et al. 2010). Indeed, immunofluorescent staining on fixed cells revealed that the branched actin marker, cortactin, localised to protrusions and to actin patches at the cell-SLB interface (Fig. 3e). Taken together this data suggests that actin polymerisation, both linear and branched, is upregulated at regions of low E-cadherin density at the cell-SLB interface.

### Branched and linear actin polymerisation are differentially regulated by cadherin mobility

Finally, to mimic the effect of changing E-cadherin mobility in a neighbouring cell-membrane on we modulated the fluidity of the SLB by changing the composition of lipids used. We prepared SLBs composed primarily of the saturated 1,2-dipalmitoyl-sn-glycero-3-phosphocholine (DPPC) that showed a 17-fold lower mobility than the otherwise used SLBs made of the unsaturated 1,2-dioleoyl-sn-glycero-3-phosphocholine (DOPC) as measured by fluorescence recovery after photobleaching of Ni-NTA tethered His-GFP (D_DOPC_ = 0.85 ± 0.05 μm^2^; D_DPPC_ = 0.05 ± 0.01 μm^2^) (Supp Fig 3 a-c, Supp Video 5). Using kymograph analysis, we found that retrograde flow is lower in cells on low mobility DPPC SLBs 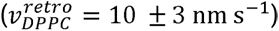 than on high mobility DOPC SLBs 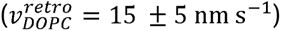 (Fig. 4a, b). To confirm that the decrease in retrograde flow is due to a decrease in Arp2/3 activity, we treated cells on DOPC SLBs with the Arp2/3 inhibitor CK-666 and found a threefold decrease in retrograde flow 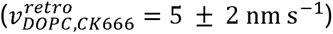. Together these results show that Arp2/3 activity is upregulated when engaging with highly mobile E-cadherins.

**Fig 4:**
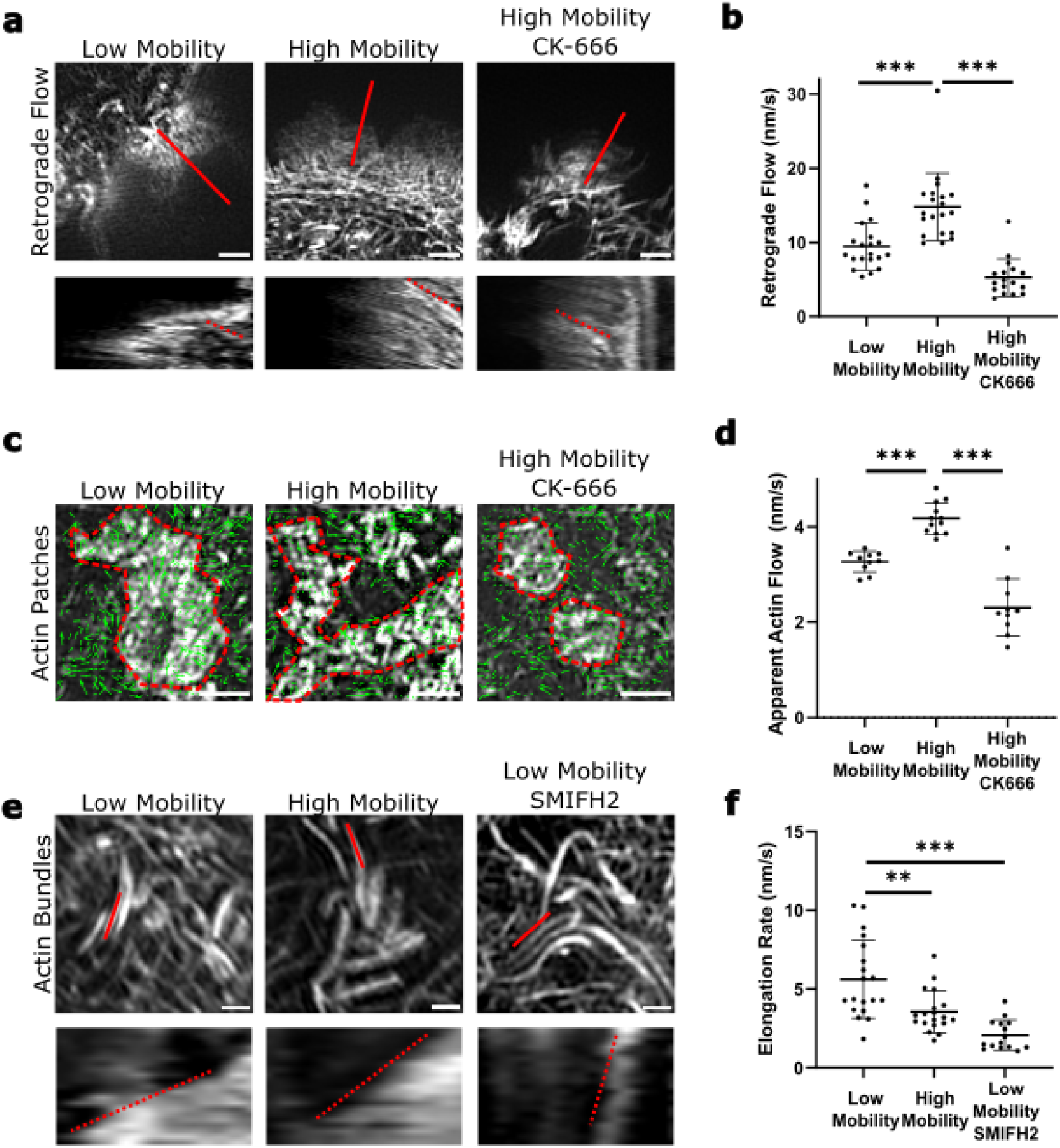
Mobility of cadherin substrate regulates branched and linear actin polymerisation. **a** Kymographs along the red line give rate of retrograde flow. Scale Bar = 1µm. **b** Rates of retrograde flow in cells plated on low and high mobility substrates and in presence of Arp2/3 inhibitor CK666. Mean ± SD plotted from N=3 **c** Velocity vectors from PIV analysis used to measure apparent actin flows in dynamic patches. Scale Bar = 1µm. **d** Apparent actin flows in dynamic patches in cells plated on low and high mobility substrates and in presence of Arp2/3 inhibitor CK666 Mean ± SD plotted from N=3. **e** Kymographs along the red line give rate of actin bundle elongation. **f** Rates of actin bundle elongation in cells plated on low and high mobility substrates and in presence of formin inhibitor SMIFH2. Mean ± SD plotted from N=3

To analyse actin flows in actin patches under the same contexts, we employed particle image velocimetry (PIV) due to the lack of directionality of actin motion (Fig. 4c). Even though the apparent actin flow velocities obtained by PIV are typically lower than the ones obtained by kymograph analysis, we could observe a 1.2 fold increase in actin flow velocity in cells on high mobility SLBs 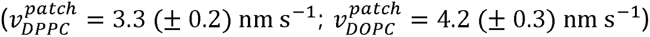 (Fig. 4d). As observed in retrograde flow, actin flow in patches decreased by a factor of 2 in cells treated with CK666 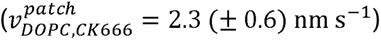, indicating that these structures are also maintained by Arp2/3-mediated actin polymerisation.

Last, we quantified actin elongation rates in actin bundles by generating kymographs along the direction of extension (Fig. 4e, Supp. Video 4). Interestingly, actin bundles showed a faster elongation rate in cells on DPPC SLBs, 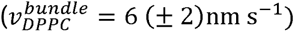 compared to DOPC SLBs 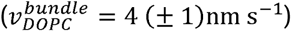, and the rate decreased threefold after cells were treated with the formin inhibitor SMIFH2 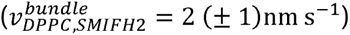 (Fig. 4f), implying that static bundles are maintained by linear actin polymerisation mediated by formins. The higher bundle elongation rate in cells on DPPC SLBs suggests that formin-mediated actin polymerisation is upregulated when E-cadherin mobility is reduced. Taken together it emerges that increased E-cadherin mobility leads to activation of Arp2/3 and a shift from linear polymerisation alone to a combination of branched and linear actin polymerisation.

To characterise how the distribution and dynamics of actin bundles and patches on a cellular level, we quantified the area occupied by actin bundles in cells where bundles and patches were present. We found that the number of actin bundles decreased within minutes (Supp Fig. 4a) and relative area they occupied (Supp Fig. 4b). Concomitantly, we found that identifiable actin fibres covered a larger relative area of E-cadherin-deficient regions at 4h compared to 5h post seeding (Supp Fig. 4c, d). We then quantified the area covered by linear and branched actin networks at different time points post-seeding (Fig 2a, Supp Fig 4e, f). After an initial abundance of branched actin at the cell periphery and disperse actin inside, actin bundles dominate as the cell-SLB interface matures. At later stages, however, regions with branched actin networks re-emerge.

## Discussion

Using super-resolution TIRF-SIM, we have shown that increase in cadherin mobility favours Arp2/3 activity while a decrease in cadherin mobility favours linear actin polymerisation by formins. We analysed temporal changes in actin architecture and found them consistent with previous reports of actin and its polymerisation factors at AJs. Early cell-cell contacts are enriched in branched actin which is reorganised into linear bundles (Lee et al. 2016; Michael et al. 2016). This switch from branched to linear actin networks is also evident from our TIRF experiments where we see a decrease in protrusion size consistent with a decrease in Arp2/3 activity after 1h post attachment. The reemergence of regions of polymerising branched actin networks at later stages could be analogous to a repair mechanism via Arp2/3 reported at late cell-cell contacts (Li et al. 2021; 2020). *In vivo*, we would expect that regions with a loss of cell-cell contact exhibit lower cadherin density due to loss of cadherin clustering and higher cadherin mobility due to detachment of AJs from the actin network. Indeed, we do see lower cadherin densities in regions of increased actin polymerisation and branched actin networks.

The actin architecture we see 1h post seeding is remarkably similar to those reported in zebrafish ectoderm progenitor cells at the same time point (Arslan et al. 2024) suggesting that the processes is conserved across cell lines and species. We also observe a flow of actin and AJs from protrusions inwards to the periphery of the cell cortex leading to the formation of the actin ring. Following the progression of this actin architecture as the cell-cell contact evolves we observed spatially heterogeneous actin polymerisation dynamics, with the appearance of dynamic actin patches and regions of seemingly stable actin bundles. We find that both dynamic patches and areas of retrograde flow behave similarly suggesting that they are maintained by the same processes. It remains unclear whether this is achieved by a recruitment and activation of Arp2/3 upon increase in cadherin mobility, for instance by the Rac1 pathway (Lee et al. 2016) or that this is a result of a release of Arp2/3 inhibition potentially inherent at mature junctions due to the presence of vinculin (James, Fokin, et al. 2025). In contrast, actin bundles are maintained by formins that are activated by the RhoA pathway as observed in integrin-mediated focal adhesion formation (Kalappurakkal et al., *Cell,* 2019). The fact that dynamic patches and static bundles appear as spatially confined structures along the cell-SLB interface and change over time suggests that RhoA and Rac1 linked pathways are confined in local zones along the cell-SLB interface (Yamada and Nelson 2007). Interestingly, the mobility of E-cadECD, and with this the amount of traction force that cells generate when dragging E-cadherin along the cell surface, could be a regulating factor for the type of actin structure generation. Global regulation of RhoA during the first hour of cell-SLB adhesion was recently reported (Arslan et al. 2024), however, our novel discovery of local zones with distinct actin patterns opens the path to investigate the biochemical and physical mechanisms allowing the cell to sense local differences in E-cadherin mobility at sub-micron scales.

## Materials and Methods

### Preparation of Supported Lipid Bilayers and Experimental Chambers

Supported lipid bilayers and the experimental chambers were prepared as described earlier(Köster et al. 2022). Glass coverslips (25 mm diameter, #1.5 borosilicate, Gerhard Menzel GmbH, cat. no. LDRND25/1.5) for SLB formation were cleaned with 2% Hellmanex III solution (Hellma Analytics, cat. No. Z805939, Merck) following the manufacturer’s instructions, thoroughly rinsed with MilliQ water, then placed in a 3M KOH (Sigma-Aldrich) solution and sonicated for 15 mins, again thoroughly rinsed with MilliQ water and finally blow dried under a stream of N2 gas.

Experimental chambers were made by cutting off the lid and conical bottom part of 0.2 ml PCR tubes (cat. no. I1402-8100, Starlab) and sticking the remaining cylinder onto the cleaned coverslip glass using UV glue (cat. no. NOA88, Norland Products) and three minutes curing by intense UV light at 265 nm (UV Stratalinker 2400, Stratagene). Freshly cleaned and assembled chambers were directly used for experiments.

Supported lipid bilayers (SLB) containing 96% of DOPC lipids (cat. no. 850375, Avanti Polar Lipids) for high mobility membranes or DPPC lipids (cat. no. 850355P, Avanti Polar Lipids) for low mobility membranes and 4% DGS-NTA(Ni2+) lipids (cat. no. 790404, Avanti Polar Lipids) were formed by fusion of small uni-lamellar vesicles (SUV) that were prepared by lipid extrusion using a membrane with 100 nm pore size (cat. no. 610000, Avanti Polar Lipids). SLBs were formed by addition of 10 µl of SUV mix at 4 mM lipid concentration to chambers filled with 90 µl SLB buffer (1xTBS solution, pH 7.4), incubated for 25 min and then washed 10 times with SLB buffer. Extrusion and SLB formation were performed at RT for the DOPC containing lipid mix and at T = 40-45nn°C for the DPPC containing lipid mix to ensure formation of uniform lipid bilayers.

Glass slides were mounted into AttoFluor metal chambers (cat. no. A7816, Invitrogen) for imaging on inverted microscopes. For live cell imaging using TIRF-SIM, no small chambers were used and SLBs were formed on the entire glass coverslip. Cleaned glass coverslips were mounted into AttoFluor chambers and SLB were formed by adding 120ul of SUV was added to 300ul SLB formation buffer.

### Cell Culture & Immunoflourescent staining

MCF 7 E-cad-GFP CRISPR-Cas9 knock in cells transduced with LifeAct-TagRFPT and iRFP670-α-catenin were a gift from Ivar Noordstra (Alpha Yap Laboratory). Cells were cultured in Dublecco’s Modified Eagle Media (DMEM) with Glutamax (Gibco) supplemented with 10% v/v fetal bovine serum (heat inactivated, cat. no. F4135, Merck), 5% (100 ug/ml) Penicillin/streptomycin, 50 ug/ml Geneticin G418 (cat. no. 10131027, Gibco) and 0.25 ug/ml Puromycin (cat. no. 15490717, Thermo Fisher) at 37 °C with 5% CO2 in a humidified incubator.

For visualising cortactin localisation, cells on SLBs were first fixed with 2% PFA for 10 mins and permeabilised with 0.1% Triton X-100 for 5 mins. Cells were stained for with primary antibodies against cortactin (4F11) and secondary antibodies labelled with Alexa405 (ab175652, Abcam) for 1h each, after 30mins blocking with 10% FBS in PBS. Fixed samples were imaged at 100x on a Nikon SoRa confocal microscope.

### Fluorescence Recovery After Photobleaching (FRAP)

We performed FRAP experiments to determine the diffusion rate of histidine-tagged proteins tethered to SLBs of different compositions. SLBs were formed in our PCR-tube based experimental chambers and labelled fluorescently by incubation with 10 nM His6-GFP for 40 min followed by washing off unbound protein (replacement of 50 ul of the old buffer with fresh one for ten times). FRAP image sequences were recorded using a spinning disc confocal microscope (Ultraview Vox, Perkin Elmer) with a 100x objective (NA 1.40, oil, Plan Apo VC, Nikon) and equipped with a FRAP unit. The sample was illuminated, and the image window was adjusted to have uniform illumination. A region of interest of 8 µm diameter was drawn using ROI tool and saved for future use. Photobleaching was performed using the 488 nm laser at maximum power. Acquisition criteria were set to 10 frames before bleaching, and 1 min (DOPC) or 4 min (DPPC) of imaging after bleaching at a time interval of 2s or 5s per frame. FRAP images were acquired with a Hamamatsu ORCA-R2 camera operated by Volocity 6.0 software (Perkin Elmer).

The mean intensity values of the FRAP region were bleach corrected using the readings of an area outside of the FRAP region and normalised by dividing the mean value from the pre-bleach image sequence. An exponential function was fitted to the recovery curve (T=0 is defined as the end of the bleach procedure), and the resulting time constant was used to calculate the diffusion constant using the equation describing fluorescence recovery of a circular region of radius 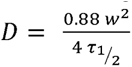 (Axelrod et al. 1976).

### Total Internal Reflection Fluorescence (TIRF) microscopy

TIRF microscopy is a method of choice to visualize fluorescently labelled objects in the vicinity of a surface while increasing the signal to noise ratio if the bulk phase also contains fluorescent species. TIRF microscopy was performed using a Nikon Eclipse Ti-E/B microscope equipped with perfect focus, a Ti-E TIRF illuminator (CW laser lines: 488 nm, 565 nm, 647 nm) and a Zyla sCMOS 4.2 camera (Andor, Oxford Instruments, UK) controlled by Andor iQ3 software. The TIRF mode was generated by illumination through the periphery of a high NA objective (100X, 1.45 NA).

### Extended-Total Internal Reflection Fluorescence – Structured Illumination Microscopy

Super-resolution imaging was performed on a custom-built TIRF-SIM setup in the Fritzsche laboratory (Oxford university) based on a ferroelectric spatial light modulator used to generate diffraction patterns and adjust the TIRF angle (Li et al. 2015). The TIRF angle was selected to ensure below 150 nm penetration depth 488, 560, and 640 nm laser lines. Illumination and detection were performed through an Olympus 100× (NA 1.49; UPLAPO100XOHR) oil objective. Raw images were obtained on two Hamamatsu Orca Flash 4.0 cameras, and reconstructed with custom made software.

### Image analysis

Cell area in TIRF images was calculated by measuring the area of actin staining utilising thresholding in FIJI. Curves were time-shifted to synchronise different fields, considering that cells were fully attached once they reach an area of 20 µm^2^. Actin, E-cadherin and α-catenin recruitment show average intensity of signal in this area normalised to the highest average intensity reached in each cell. To calculate protrusion area, cell edges were first segmented using Otsu thresholding and a morphological opening with a 2.68μm filter was applied to generate a smoothened inner edge corresponding to the cell cortex. Protrusion area was then calculated as the area between the detected and smoothed edges.

Rate of photodamage is the rate of decrease in relative actin intensities in manually drawn ROIs containing different actin architectures, normalised to the intensity at the first time-point. Rate of growth of static bundles and rate of retrograde flow were measured from kymographs drawn along the direction of growth and flow respectively. PIV was carried out using PIVLAB in MATLAB. Each point on the graph represents average magnitudes of velocities at all points within a region of dynamic actin, further averaged over 30 time intervals. Areas of regions containing different actin architectures were estimated by manually drawing ROI in FIJI. For detecting actin bundles, linear structures within manually drawn ROIs were detected using the Sato tubeness enhancement filter. Actin bundles were then selected after segmentation, by manually setting a threshold at the first time point. Threshold values for subsequent time points were normalised against bleaching across the whole field.

For statistical analysis, populations were first tested for normality using a Shapiro-Wilk test. Difference between means was tested using t-test with Welch’s correction for populations conforming to normality and with Mann-Whitney test for population deviating from normality.

## Acknowledgements

The authors thank CAMDU (Computing and Advanced Microscopy Development Unit) at University of Warwick and Yu Zhang and Dr. Jacky (Ka Long) Ko at the Kennedy Institute for Rheumatology, University of Oxford, for their support & assistance in this work. SG, JJ, BHU and DK would like to thank our colleagues at the Centre for Mechanochemical Cell Biology for inspiring conversations and creating a collaborative work environment. SG and BHU were supported by the University of Warwick International Chancellor’s fellowship, JJ and DK were supported by grants from EPSRC (EP/V043498/1; EP/Y002245/1), and SG and DK gratefully acknowledge funding from the Medical and Life Sciences Research Fund. MF acknowledges funding from MRC (APP23835) and CRUK (EP/X033015/1).

## Figures

**Supp Fig 1:**
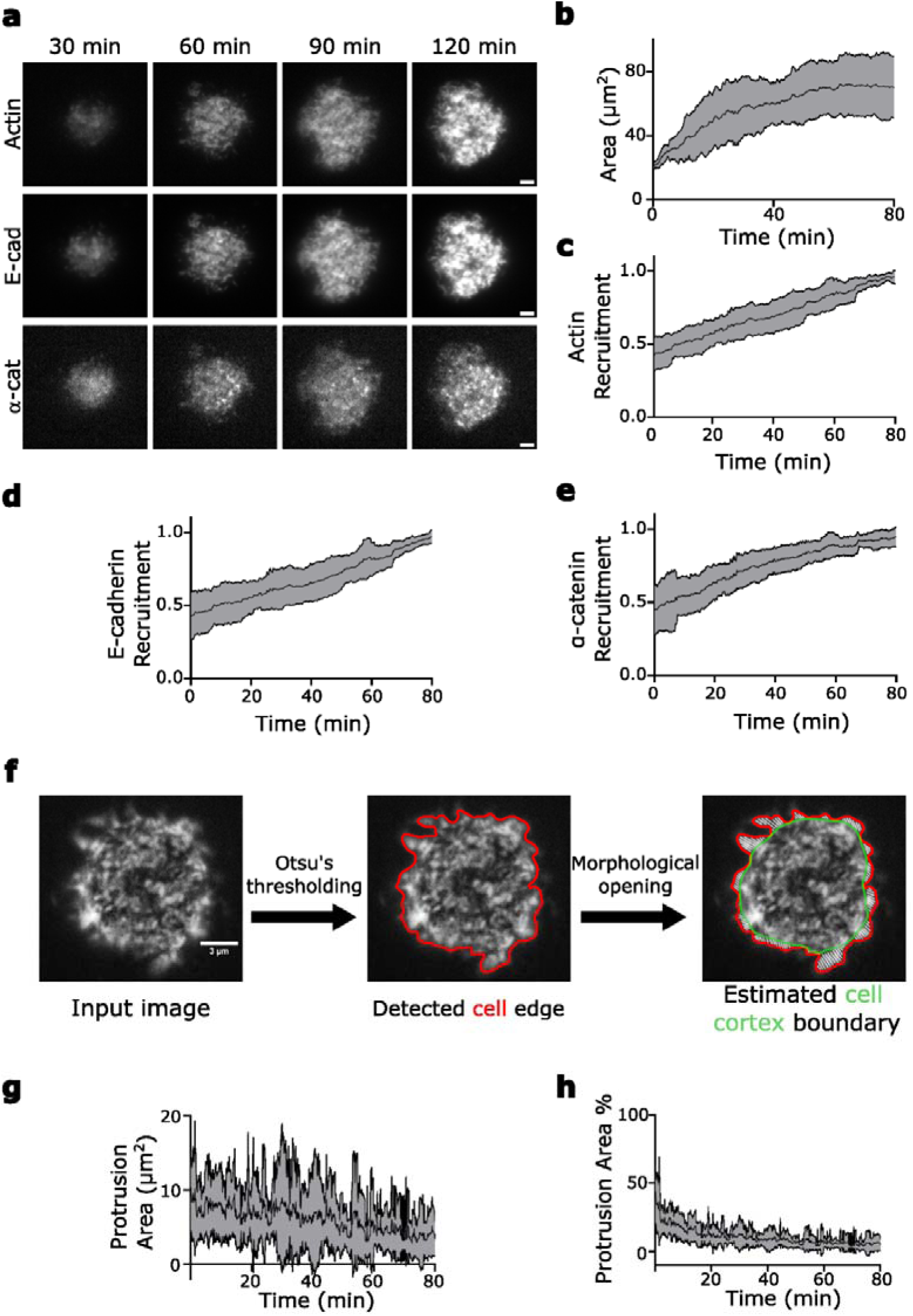
**a** TIRF images of MCF7 cells with E-cadherin-GFP knock in cells, stably expressing LifeAct-TagRFPT and iRFP670-α-catenin plated on SLBs decorated with His_6_-E-cadECD Scale Bar = 2µm. **b** Area of individual cells spreading over time. Mean ± SD plotted n=7. **c-e** Plots of relative intensities of LifeAct-TagRFPT, GFP-E-cadherin, and iRFP670-α-catenin at the cell-SLB interface over time. Mean ± SD of n=14 **f** Protrusion area is calculated as the area between detected edge of the cell (Red) and the estimated edge of the cortex (Green). **g,h** Protrusion area expressed in µm^2^ and as a percentage of cell area. Mean ± SD plotted n=7.

**Supp Fig 2:**
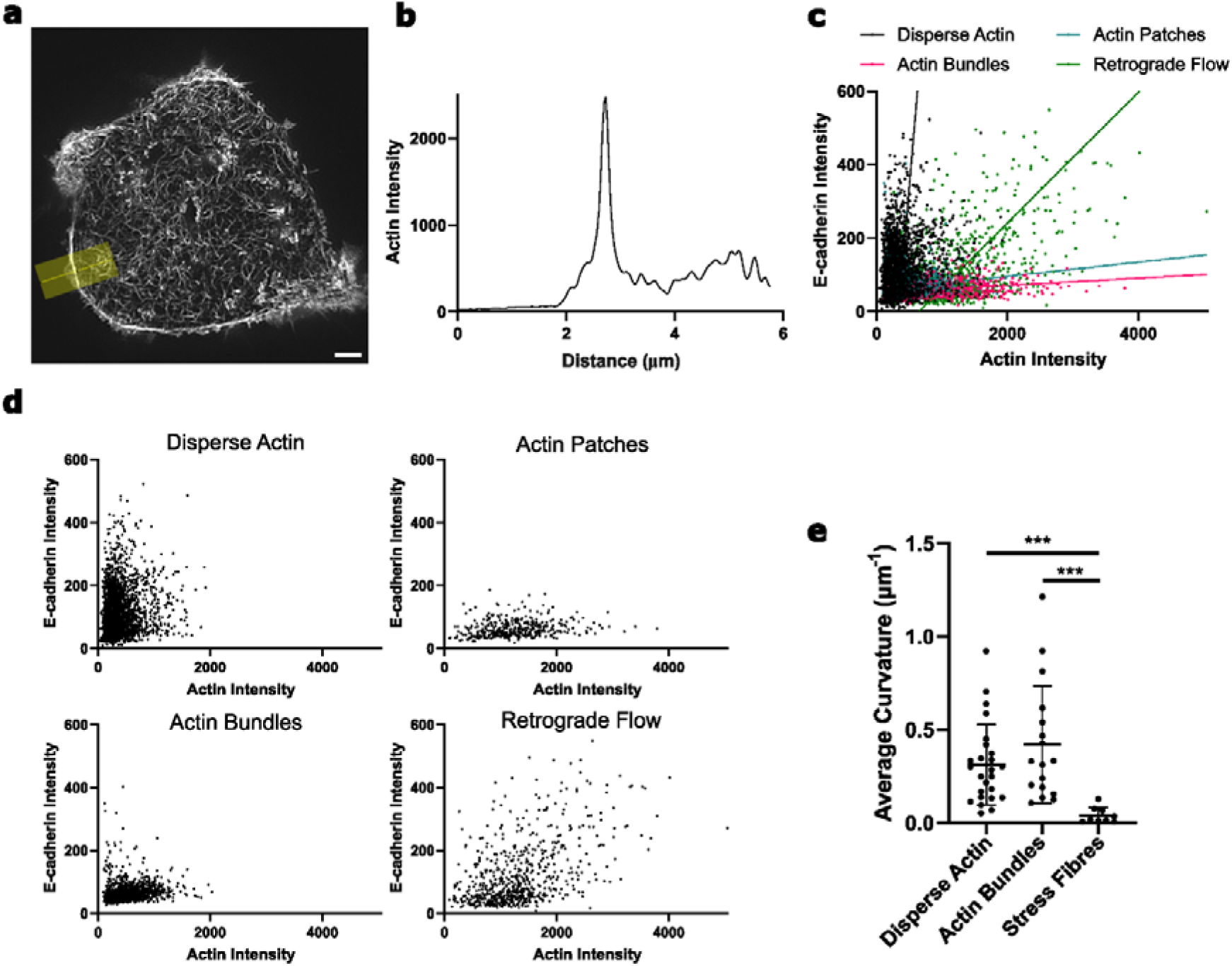
**a** TIRF-SIM image of an MCF7 cell expressing LifeAct-TagRFPT, showing formation of actin ring. **b** Line scan of actin intensity along region shown in **a. c,d** Intensities of LifeAct-TagRFPT and GFP-E-cadherin in regions of different actin architecture. Deming regression lines show relationship between actin and E-cadherin intensities in different actin architectures. **e** Average curvatures of disperse filaments and actin bundles in MCF7 cells plates on E-cadECD SLBs along with stress fibres in MCF7 cells plated on fibronectin-coated glass coverslips.

**Supp Fig 3:**
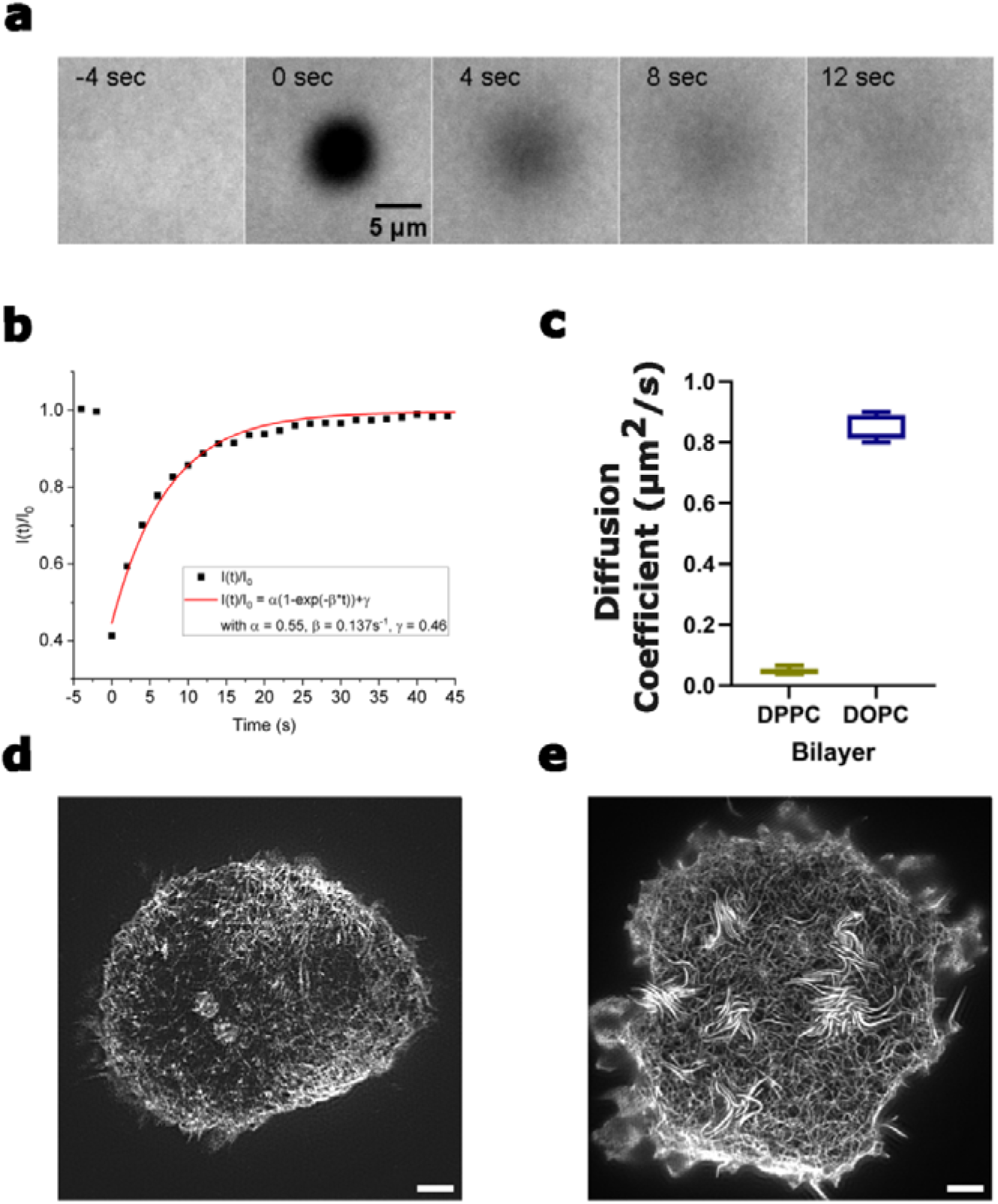
**a** TIRF Images of DOPC SLBs decorated with EGFP showing fluorescence recovery after photobleaching (FRAP). **b** Intensity plot of fluorescence recovery over time after photobleaching used to calculate diffusion coefficient of the SLB. **c** Diffusion coefficient of DPPC and DOPC SLBs. Mean ± SD plotted. **d** Sample image of whole cell for triple-labelled MCF10A cells plated on DOPC after treatment with 100µM CK666. **e** Sample image of whole cell for triple-labelled MCF10A cell plated on DPPC after treatment with SMIFH2.

**Supp Fig 4:**
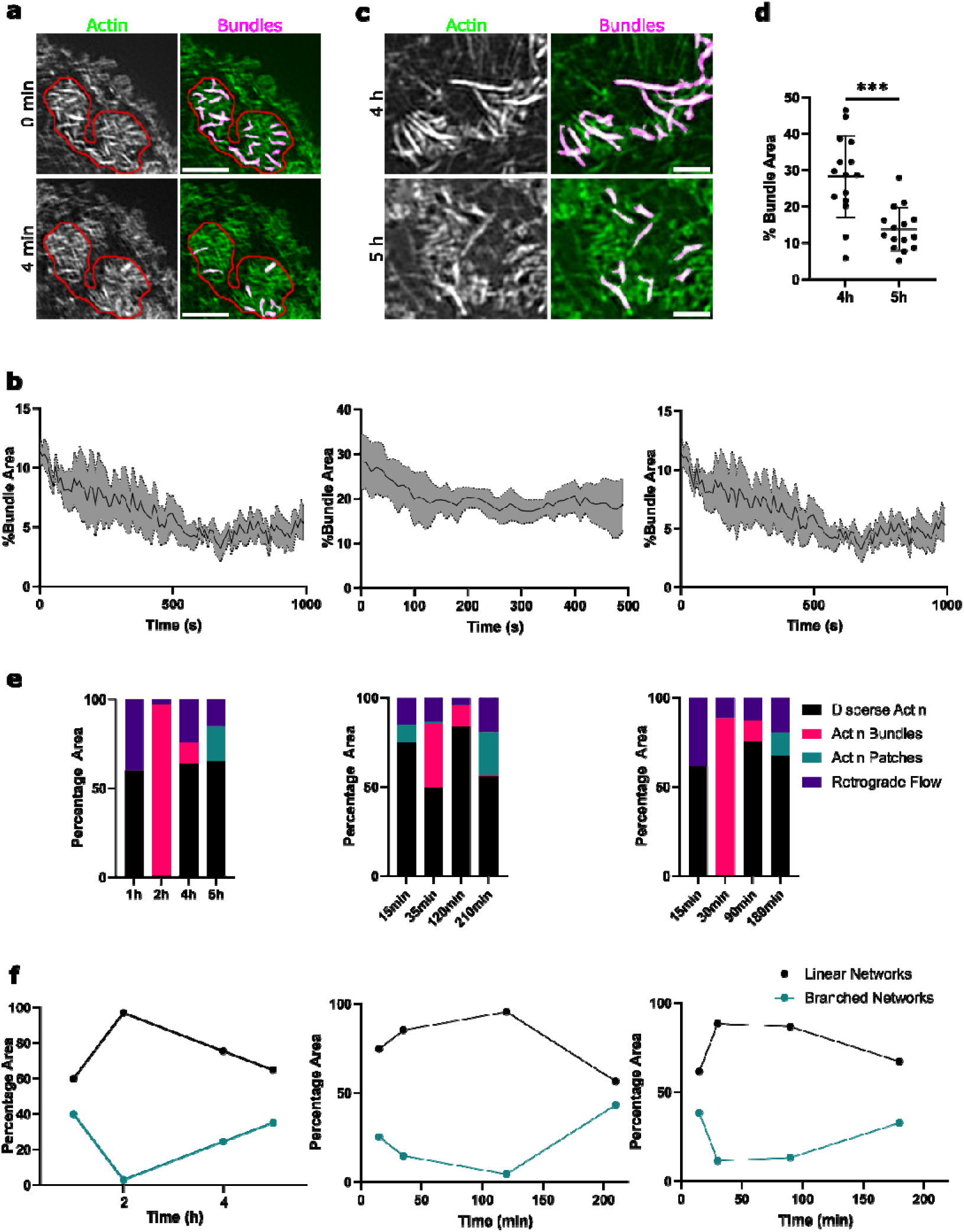
**a** TIRF-SIM images of actin and detected bundles within ROIs imaged over short time periods. Scale Bar = 2µm **b** Plots showing decrease in % bundle area over time in 3 independent repeats. Mean ± SD plotted **c** TIRF-SIM images of actin and detected bundles 4 and 5 h post seeding **d** % Bundle Area in E-cadherin deficient patches 4 and 5 hours post-seeding. Mean ± SD plotted. N=3 **e** Plots showing percentage area of the cell covered by different actin architectures at different times post seeding in 3 independent repeats. **f** Plots showing percentage area of the cell covered by branched and linear actin networks at different times post seeding in 3 independent repeats.

